# Environmental Assessment of Cytotoxic Drugs in the Oncology Center of Cyprus

**DOI:** 10.1101/610931

**Authors:** Elpidoforos S. Soteriades, Sofia C. Economidou, Artemis Tsivitanidou, Petros Polyviou, Amanda Lorimer, Nikos Katodritis, Stavroula Theophanous-Kitiri

## Abstract

**Background:** Cytotoxic drugs constitute an important workplace hazard in the hospital environment. Our aim was to conduct an environmental assessment of hazardous drugs in the Oncology center of Cyprus.

**Methods:** Wipe samples were obtained from 42 workplace areas of the Oncology Center including two pairs of gloves in an initial assessment, while 10 samples were obtained at follow-up 3 years later. Potential contamination with cyclophosphamide (CP), ifosphamide (IF) and 5-fluorouracil (5-FU) and other cytotoxic medications was examined using the GC-MSMS system (CP, IF) and the HPLC system with UV detection (5-FU) method, respectively.

**Results:** Wipe sample contamination was detected at 11.9% and 15% in the initial and follow-up assessment, respectively. Both pairs of gloves assessed were free from contamination. The results showed contamination with cyclophosphamide on the work space inside the isolator, on a day-care office phone and on the central pharmacy bench. Ifosphamide was only detected on the floor of a patient’s room. Contamination with 5-fluorouracil was found only on the surface of a prepared IV infusion bag. The levels of contamination in the positive samples ranged from 0.05 to 10.12 ng/cm^2^.

**Conclusions:** The overall percentage of sample contamination at the Oncology Center was very low compared to other centers around the world. In addition, the detected levels of contamination with cytotoxic drugs were relatively low with the exception of the workspace inside the biological safety cabinet.

These results in both assessments may reflect the implementation of comprehensive control measures including employee training, technological equipment and effective cleaning procedures.

## Introduction

Anti-neoplastic agents constitute a significant workplace hazard for health professionals in the hospital environment [1, 2]. Such hazardous drugs, used in the treatment of cancer, have been associated with many adverse health effects following employee acute and/or chronic cumulative exposure. Cytotoxic drugs have been particularly associated with reproductive toxicity as documented by several scientific publications in the international literature [3–5]. However, such reported reproductive toxicity was linked to higher levels of workers’ exposure to such drugs, usually observed in past decades [usually at the level of milligrams per milliliter (mg/mL) or (mg/cm^2^)] compared to current levels of potential exposure observed nowadays [nanograms per milliliter (ng/mL) or (ng/cm^2^)] [6–8].

Environmental and biological monitoring has been extensively used in order to detect potential workplace exposure of health professionals to anti-neoplastic agents and evaluate associated health risks [9–11]. Several methods have been employed to measure the level of environmental contamination with hazardous drugs in relevant occupational settings [12–15]. Collecting surface wipe samples from different sites of the hospital environment constitutes one of the most widespread method used for environmental assessment studies around the world [16–18]. Sugiura et al reported multi-center environmental monitoring studies evaluating cyclophosphamide exposure in Japan with levels of contamination ranging from 50% to 80% among all samples collected [19, 20]. In a study conducted in 6 British Columbian hospital pharmacies, 61% of the samples tested positive for contamination with Cyclophosphamide (CP) or Methotrexate (MTX). It is worth mentioning that contamination was detected sometimes even after cleaning, implying that the cleaning protocols in British Colombia hospitals required further improvement [16]. In Sweden, a similar study in a hospital pharmacy showed that CP or Ifosfamide (IF) were detected and quantified in 96-100% of wipe samples obtained from several areas in the preparation unit, with the highest values observed in the dressing room. However, after the cleaning procedures were reviewed and a second measurement was conducted several months later with samples taken from the same sites, results showed significantly lower levels of contamination [17]. Another study performed in two hospitals in France, where positive air pressure isolators were used, demonstrated much lower levels of contamination especially in areas outside the isolators [21].

In the US, between 2000–2005, the closed-system drug transfer device (CSTD) was introduced, and a study of environmental assessment was conducted in 22 hospitals. When standard drug delivery devices were used, the levels of contamination with positive wipe samples were 78%, 54% and 33% for Cyclophosphamide, Ifosfamide and 5-fluorouracil, respectively. With the introduction of the closed-system drug transfer device, there was a significant decrease in the levels of contamination to 68%, 45% and 20%, respectively [22]. Several other studies have shown significant reduction in the levels of occupational and environmental contamination as well as exposure to hazardous drugs following the implementation of a CSDT device [23–27].

In Cyprus, there has been no previous study evaluating the potential contamination of hospital environments with hazardous drugs in the oncology units. The objective of our study was to assess the potential workplace contamination of the main oncology center of Cyprus with three most frequently used cytotoxic drugs, namely Cyclophospamide, Ifosphamide and 5-fluorouracil. In addition, we conducted a follow-up assessment study evaluating potential contamination with a number of other cytotoxic drugs three years later.

## Methods

### Hospital setting

An environmental contamination assessment was conducted at the Bank of Cyprus Oncology Center (BOCOC) in 2011 and a follow-up assessment was repeated in 2014 in the context of a European comparative study between different hospitals. BOCOC is the main oncology center of the island providing care to about 70% of the cancer patients in Cyprus. The Center has 32 beds, has a workforce of about 200 employees, and provides outpatient cancer treatment to about 60 patients on a daily basis. It has two inpatient wards, a central pharmacy and an outpatient day care facility. Preparation of cytotoxic drugs is performed by trained nurses in a specifically designed unit equipped with two biological safety cabinets that are externally vented. The cleaning/decontamination protocols of our center includes daily cleaning of the biological safety cabinets. The center is required to prepare about 70 separate treatment protocols on a daily basis for the needs of in-patients and the day-care therapy unit. The units of Cytotoxic treatments are administered by nurses to patients in the wards and also in a daycare unit. Between the two environmental assessments we were able to introduce the use of a closed system drug transfer device (CSTD) in our Center.

### Wipe sample collection

Wipe samples were taken from 42 workplace surfaces including two pairs of gloves for the initial assessment, while in the follow-up assessment, a total of 10 samples were obtained. The initial wipe samples were taken with Cyto Wipe Kits from Exposure Control Sweden AB [27]. Samples were obtained from all departments of the oncology center (central pharmacy, outpatient pharmacy, chemotherapy pharmacy, day care unit, patient wards, radiotherapy department and administration offices). In November 2011, wipe samples were taken and gloves were collected by a nurse under the supervision of the head of the pharmacy department. The contamination in nanograms per cm^2^ was calculated assuming 100% recovery and wipe efficiency. This means that all results may be considered as underestimates. The detection limits for the analysis of cyclophosphamide, ifosphamide and 5-fluorouracil were 0.10, 0.10 and 5 ng/mL extract, respectively. The methodology used for the collection of the follow-up samples in 2014 was quite similar although the samples were sent to a different laboratory. In addition, the same health professionals were involved in the sample collection as in the initial assessment. The second assessment was performed in the context of a European comparative study between different hospitals in Europe. A permission to use the results for our Center from the second study has been obtained. Although fewer samples were used in the second assessment, the samples were taken from similar sites as the initial assessment for the purpose of comparison over time. The sites from each wipe study are described in detail in each table (Tables 1 and 2). The follow-up samples were sent to the Institute of Energy and Environmental Technology in Germany (IUTA). Follow-up analyses were performed for the following hazardous drugs: 5-fluorouracil, Gemcitabine, Methotrexate, Topotecan, Irinotecan, Doxorubicin, Epirubicin, Ifosfamide, Cyclophospamide, Etoposide, Docetaxel, Paclitaxel with the corresponding detection limits: 0.008, 0.003, 0.003, 0.003, 0.003, 0.003, 0.003, 0.008, 0.008, 0.008, 0.02, 0.02 ng/cm^2^, respectively. There was no change in environmental controls (hoods, rooms etc.) from the first wipe to the second wipe study.

**Table 1:**
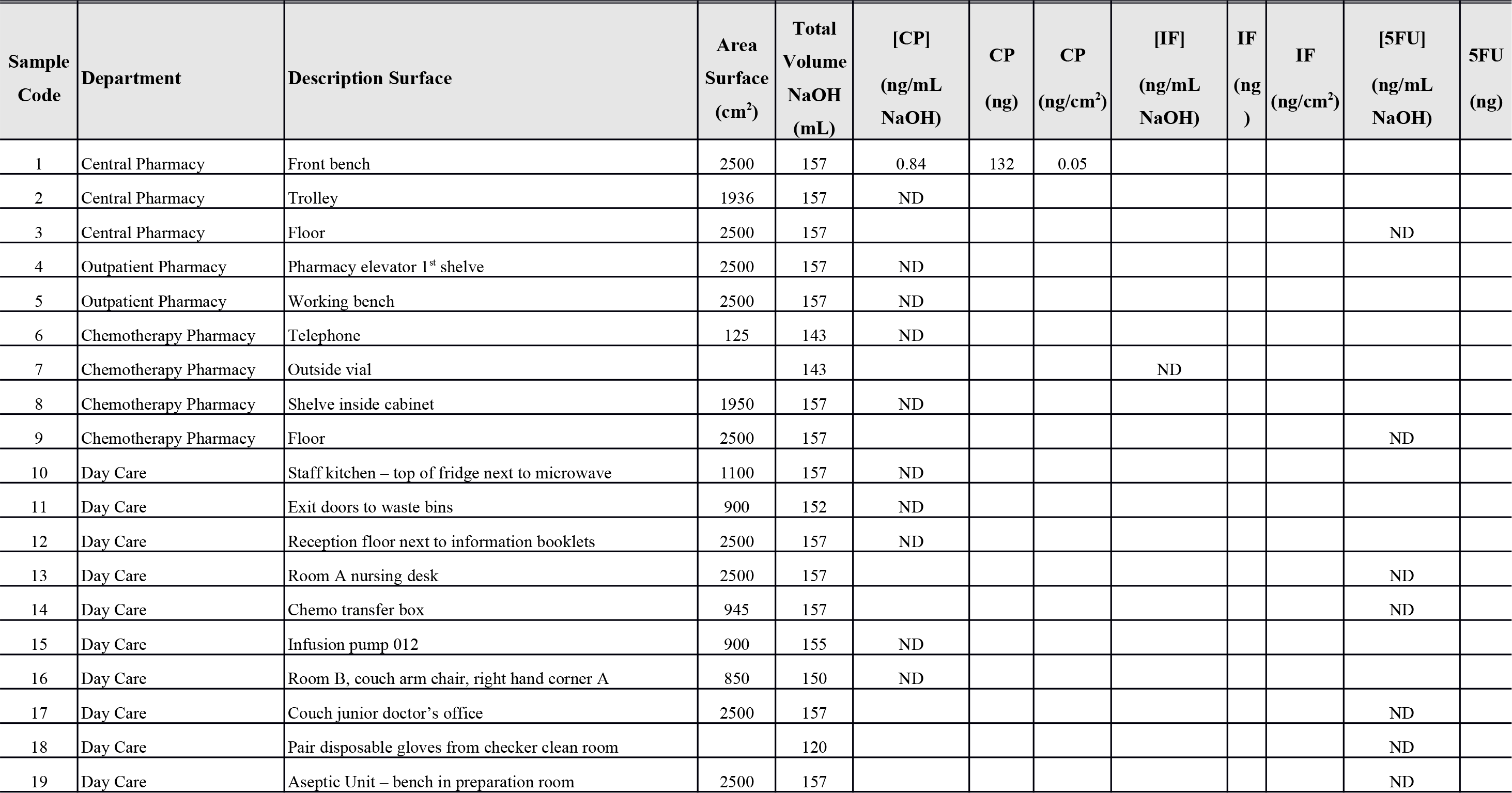
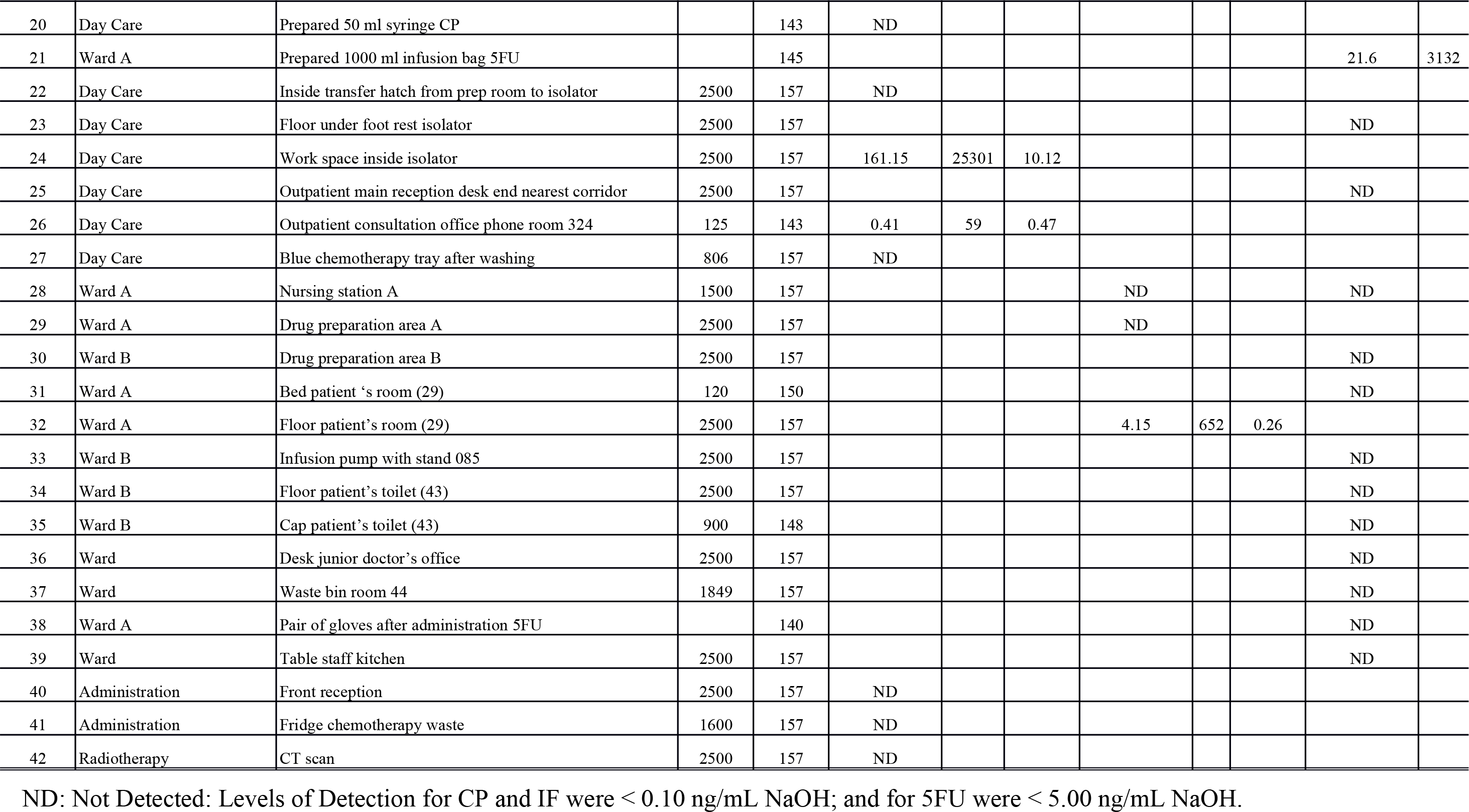
Results of a wipe-sample environmental assessment conducted in 2011 evaluating potential contamination with Cyclophosphamide (CP), ifosphamide (IF) and 5-fluorouracil (5FU) at the Bank of Cyprus Oncology Center – Initial assessment

| Sample Code | Department | Description Surface | Area Surface (cm<sup>2</sup>) | Total Volume NaOH (mL) | [CP] (ng/mL NaOH) | CP (ng) | CP (ng/cm<sup>2</sup>) | [IF] (ng/mL NaOH) | IF (ng) | IF (ng/cm<sup>2</sup>) | [5FU] (ng/mL NaOH) | 5FU (ng) |
| --- | --- | --- | --- | --- | --- | --- | --- | --- | --- | --- | --- | --- |
| 1 | Central Pharmacy | Front bench | 2500 | 157 | 0.84 | 132 | 0.05 |  |  |  |  |  |
| 2 | Central Pharmacy | Trolley | 1936 | 157 | ND |  |  |  |  |  |  |  |
| 3 | Central Pharmacy | Floor | 2500 | 157 |  |  |  |  |  |  | ND |  |
| 4 | Outpatient Pharmacy | Pharmacy elevator 1<sup>st</sup> shelve | 2500 | 157 | ND |  |  |  |  |  |  |  |
| 5 | Outpatient Pharmacy | Working bench | 2500 | 157 | ND |  |  |  |  |  |  |  |
| 6 | Chemotherapy Pharmacy | Telephone | 125 | 143 | ND |  |  |  |  |  |  |  |
| 7 | Chemotherapy Pharmacy | Outside vial |  | 143 |  |  |  | ND |  |  |  |  |
| 8 | Chemotherapy Pharmacy | Shelve inside cabinet | 1950 | 157 | ND |  |  |  |  |  |  |  |
| 9 | Chemotherapy Pharmacy | Floor | 2500 | 157 |  |  |  |  |  |  | ND |  |
| 10 | Day Care | Staff kitchen – top of fridge next to microwave | 1100 | 157 | ND |  |  |  |  |  |  |  |
| 11 | Day Care | Exit doors to waste bins | 900 | 152 | ND |  |  |  |  |  |  |  |
| 12 | Day Care | Reception floor next to information booklets | 2500 | 157 | ND |  |  |  |  |  |  |  |
| 13 | Day Care | Room A nursing desk | 2500 | 157 |  |  |  |  |  |  | ND |  |
| 14 | Day Care | Chemo transfer box | 945 | 157 |  |  |  |  |  |  | ND |  |
| 15 | Day Care | Infusion pump 012 | 900 | 155 | ND |  |  |  |  |  |  |  |
| 16 | Day Care | Room B, couch arm chair, right hand corner A | 850 | 150 | ND |  |  |  |  |  |  |  |
| 17 | Day Care | Couch junior doctor's office | 2500 | 157 |  |  |  |  |  |  | ND |  |
| 18 | Day Care | Pair disposable gloves from checker clean room |  | 120 |  |  |  |  |  |  | ND |  |
| 19 | Day Care | Aseptic Unit – bench in preparation room | 2500 | 157 |  |  |  |  |  |  | ND |  |
| 20 | Day Care | Prepared 50 ml syringe CP |  | 143 | ND |  |  |  |  |  |  |  |
| 21 | Ward A | Prepared 1000 ml infusion bag 5FU |  | 145 |  |  |  |  |  |  | 21.6 | 3132 |
| 22 | Day Care | Inside transfer hatch from prep room to isolator | 2500 | 157 | ND |  |  |  |  |  |  |  |
| 23 | Day Care | Floor under foot rest isolator | 2500 | 157 |  |  |  |  |  |  | ND |  |
| 24 | Day Care | Work space inside isolator | 2500 | 157 | 161.15 | 25301 | 10.12 |  |  |  |  |  |
| 25 | Day Care | Outpatient main reception desk end nearest corridor | 2500 | 157 |  |  |  |  |  |  | ND |  |
| 26 | Day Care | Outpatient consultation office phone room 324 | 125 | 143 | 0.41 | 59 | 0.47 |  |  |  |  |  |
| 27 | Day Care | Blue chemotherapy tray after washing | 806 | 157 | ND |  |  |  |  |  |  |  |
| 28 | Ward A | Nursing station A | 1500 | 157 |  |  |  | ND |  |  | ND |  |
| 29 | Ward A | Drug preparation area A | 2500 | 157 |  |  |  | ND |  |  |  |  |
| 30 | Ward B | Drug preparation area B | 2500 | 157 |  |  |  |  |  |  | ND |  |
| 31 | Ward A | Bed patient 's room (29) | 120 | 150 |  |  |  |  |  |  | ND |  |
| 32 | Ward A | Floor patient's room (29) | 2500 | 157 |  |  |  | 4.15 | 652 | 0.26 |  |  |
| 33 | Ward B | Infusion pump with stand 085 | 2500 | 157 |  |  |  |  |  |  | ND |  |
| 34 | Ward B | Floor patient's toilet (43) | 2500 | 157 |  |  |  |  |  |  | ND |  |
| 35 | Ward B | Cap patient's toilet (43) | 900 | 148 |  |  |  |  |  |  | ND |  |
| 36 | Ward | Desk junior doctor's office | 2500 | 157 |  |  |  |  |  |  | ND |  |
| 37 | Ward | Waste bin room 44 | 1849 | 157 |  |  |  |  |  |  | ND |  |
| 38 | Ward A | Pair of gloves after administration 5FU |  | 140 |  |  |  |  |  |  | ND |  |
| 39 | Ward | Table staff kitchen | 2500 | 157 |  |  |  |  |  |  | ND |  |
| 40 | Administration | Front reception | 2500 | 157 | ND |  |  |  |  |  |  |  |
| 41 | Administration | Fridge chemotherapy waste | 1600 | 157 | ND |  |  |  |  |  |  |  |
| 42 | Radiotherapy | CT scan | 2500 | 157 | ND |  |  |  |  |  |  |  |

ND: Not Detected: Levels of Detection for CP and IF were < 0.10 ng/mL NaOH; and for 5FU were < 5.00 ng/mL NaOH.

**Table 2:**
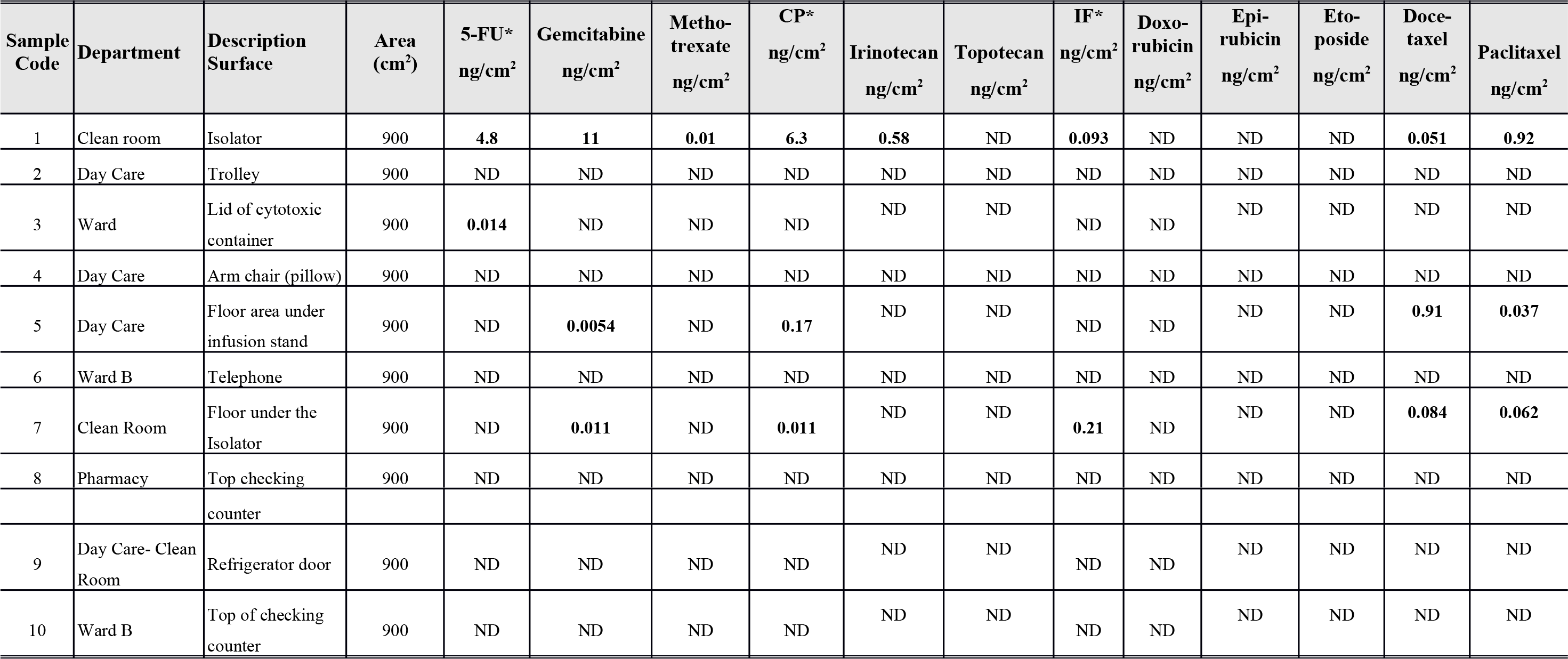
Results of a wipe-sample environmental assessment conducted in 2014 evaluating potential contamination with 10 cytotoxic drugs at the Bank of Cyprus Oncology Center - Follow up assessment

### Storage, transportation and analysis of samples

On the initial assessment, all samples were stored frozen after sampling and during transport until sample preparation and analysis. The wipe samples were prepared by adding 140 mL of a 0.03 M NaOH solution. For the gloves, 120 or 140 mL solution was used. After extraction, a part of the extract was further cleaned up according to standard procedures [28, 29]. All samples were frozen right after collection and were sent to Exposure Control B.V, the Netherlands where analysis was performed. Cyclophosphamide and ifosphamide were analysed using a Gas Chromatography – Mass Spectrometry (GC-MS) method on a GC-MSMS system showing increased sensitivity and specificity [30–31]. The analysis of 5-fluorouracil was performed on a High Performance Liquid Chromatography (HPLC) system with UV detection. The second environmental assessment was also performed using LC-MS/MS. The results were reported in nanograms per centimeters squared (ng/cm^2^).

## Results

The results of the initial environmental assessment analyses of the wipe samples are presented in Table 1. The study findings on the initial assessment show contamination with cyclophosphamide on the work surface inside the isolator as expected (biological safety cabinet), on an office phone at the daycare unit, and on the front of the bench in the central pharmacy. Ifosphamide was only observed on the floor of a patient’s room at Ward A. Contamination with 5-fluorouracil was not found in the environment. A diluted IV bag was contaminated with 5-fluorouracil on the outside surface of the bag. Except for the work space inside the isolator and the IV bag, the levels of contamination were very low. The two pair of gloves examined were not found to be contaminated with 5-fluorouracil. Overall, only 11.9% of the samples were tested positive for any of the three cytotoxics drugs evaluated.

The results of the follow-up assessment are presented in Table 2. The follow-up assessment included a total of 12 different medications on 10 environmental samples. Based on the total number of samples multiplied by the total number of drugs tested, we found 18 positive samples with a percentage of contamination at 15% of the total samples tested. The Lid of cytotoxic waste container tested positive for contamination with 5-FU, while the floor area under the infusion stand at the daycare unit tested positive for Gemcitabine, Cyclophospamide, Docetaxel and Paclitaxel.

## Discussion

The results of both environmental assessments showed quite low and similar levels of contamination in the Oncology Center of Cyprus. In the initial assessment, the contamination with cyclophosphamide or ifosphamide was detected mainly at the daycare unit and to a lesser extent in the central pharmacy and patient wards. Spread of contamination was not observed. The contamination with cyclophosphamide in the space inside the isolator, was expected to be relatively high, however the levels of contamination on the other positive samples were very low. Contamination with 5-fluorouracil was not found in the environment. This may probably be attributed to a higher detection limit for the analysis of 5-fluorouracil compared to cyclophosphamide and ifosphamide. Overall the percentage of positive samples was much lower compared to other international scientific reports [16-20, 28, 29, 32, 33]. Low levels of contamination were also observed on the follow-up assessment conducted three years later. The difference between the two assessments is not significant (12% vs 15%) and could most likely attributed to the small number of samples used in the second assessment and the much bigger number of drugs implicated in the testing. We believe that the two assessments in general provide a similar picture of the low level contamination in our center. Except from the contamination detected in the clean room isolator and the floor underneath it, which was expected, only two other samples tested positive (the lid of cytotoxic waste container and the floor area under the infusion stand at the daycare). A sample obtained from a telephone on Ward B was negative for contamination on the follow-up assessment, compared to the wipe sample from a telephone in an office room that tested positive for contamination on the first assessment. The initial result was most likely related to unsafe practices of health professionals.

We believe that the low overall contamination levels with hazardous drugs seen in our study, may partly be attributed to the relatively small size of the Oncology Center where employees interact and cooperate within a somewhat family environment and are being concerned not only for their own health but also for their colleagues. It could also be due to the periodic repeated training of nurses and pharmacists in our Center on cytotoxics’ management based on quality accreditation requirements, and the close/daily supervision of cleaning practices. The improvement seen in the second assessment was most likely associated with additional training provided. The training consisted of two one-hour presentations from the head of pharmacy and the Occupational physician of the Center and was delivered to all nurses, pharmacists, and junior physicians of the Center. It concluded with a question/answer session.

The observed environmental contamination indicates several potential sources. A well-documented source of contamination is associated with spillage during preparation and administration of the cytotoxic drugs inside the isolator. This is also supported by the contamination found on the prepared IV infusion bag in the initial assessment and the floor area under the infusion stand in the follow-up study. In addition, another source of potential contamination includes the fact that the external vials may also be contaminated [34–36]. Although somewhat expected, contamination inside the isolator and on the pharmacy bench can easily be transferred to prepared IV bags and further spread into the hospital environment. A similar limited spread was the observed contamination of the medical office phone found in the first assessment. Such findings support the need for health professionals to use gloves whenever they come into contact with prepared IV bags and other potential contaminated products, surfaces and/or vials associated with hazardous drugs. In addition, comprehensive cleaning protocols are essential. The rather high level of contamination on the workspace inside the isolator at the daycare unit and on the prepared IV infusion bag requires further investigation. Prevention and/or control of spillages in the isolator and the IV bags may be achieved through continuous employee training and comprehensive cleaning procedures, which would limit consequent spread of contamination with cytotoxic drugs in the hospital environment and lead to minimal and/or no subsequent exposure of health professionals to such cytotoxic drugs [8].

Some limitations of our study need to be acknowledged. Although this is the first study conducted in an oncology hospital in Cyprus, we lack reference data from other local hospitals in order to perform useful comparisons. In addition, due to cost limitations, we have used a relatively small number of environmental samples however we believe that the number is sufficient to obtain a comprehensive evaluation given the relatively small size of the Oncology Center. Our study was only focused on the environmental assessment for cytotoxic drugs. Nevertheless, it could have benefited from a parallel biological monitoring study to assess potential occupational exposure among health professionals.

In conclusion, this is the first study conducted in Cyprus examining the potential environmental contamination with cytotoxic drugs in an oncology hospital setting. Our results show a relatively low level of contamination compared to a number of similar studies around the world. The outcome of our study in both assessments as well as the relatively low levels of environmental contamination with hazardous drugs could be attributed to a number of workplace practices including the use of a specifically designed isolator unit with biological safety cabinets externally vented through a separate ventilation system for the whole isolator unit. Furthermore, the employees involved in the drug preparation processes are highly trained and also receive annual refresher courses. The maintenance of a continuous monitoring system for dilution procedures run by trained pharmacy personnel could also play a role along with the adoption of the European Oncology Pharmacy protocol for monitoring spillages in the workplace. In addition to the above measure, between the two assessments, we had introduced the closed system drug transfer device (CSTD) in our Center. Finally, we believe that the cleaning protocols followed including a daily routine cleaning of the isolator system are contributing to the low contamination levels found. The slightly higher percentage of the positive samples in the second assessment may reflect the smaller number of samples obtained and the much larger number of medications tested. We believe that the above combination of workplace control measures may constitute a good practice that could be followed by other centers around the world in order to lower potential contamination levels and reduce associated exposure of health professionals to hazardous drugs in the workplace.

